# Candidozyma auris utilizes transferrin, but not heme-bound iron for in vivo virulence

**DOI:** 10.64898/2026.06.09.731159

**Authors:** Tanmay Arekar, Divya Katikaneni, Thrisha Acharya, Katherine Horst, Guolei Zhao, Gabriella Garcia, Sofia Hernalsteen, Chelsea K. Weber, Abhishek Gour, Tracy Punshnon, Abhisheak Sharma, Michail S. Lionakis, Teresa R. O’Meara, Yogesh Scindia

## Abstract

*Candidozyma auris (C. auris)* is an emerging multidrug-resistant fungal pathogen, and its dissemination to the bloodstream and deep-seated organs is associated with high mortality. The limited antifungal armory and pipelines against *C. auris* pose a major challenge in disease management. Addressing this threat requires a deeper understanding of fungal virulence mechanisms that promote persistence and of host factors that drive susceptibility. Previous in vitro studies showed that iron enhances *C. auris* resistance to azoles and echinocandins, whereas iron chelation mitigates this effect. Here, we demonstrate that *C. auris* does not utilize cell-free heme or induce hemolysis but instead extracts and uses iron from transferrin to support growth and virulence. Deletion of the *SIT1* siderophore transporter in *C. auris* attenuated fungal growth and reduced renal injury, while increased transferrin-iron saturation worsened disease outcomes in immunocompetent mice, highlighting the importance of transferrin-bound iron uptake. Mechanistically, *C. auris* exploits transferrin-bound iron to enhance ergosterol biosynthesis and activate antioxidant defenses, promoting resistance to neutrophil- and caspofungin-mediated killing. These findings identify elevated transferrin saturation as a novel host susceptibility risk factor for disseminated *C. auris* infection and reveal how iron availability reshapes fungal physiology to drive infection persistence.

## INTRODUCTION

*Candidozyma auris (C. auris)* has been designated as a critical-priority fungal pathogen due to its rapid global emergence, multidrug resistance, and high mortality rate(*1, 2*). Approximately 25% of colonized individuals progress to candidemia within 60 days(*3*), with mortality rates ranging from 40% to 67%(*4, 5*). Resistance can also emerge during therapy and fuel hospital outbreaks, underscoring the limited effectiveness of current antifungal-centered strategies(*6*). While we continue to advance our understanding of *C. auris* biology and the factors influencing its transition to a virulent state, our appreciation of host-related risk factors driving susceptibility to infection remains limited. Thus, there is an unmet medical need to define host mechanisms that govern susceptibility, fungal persistence and treatment failure, and to develop novel non-conventional preventive strategies for high-risk patients.

Iron is essential for host physiology(*7, 8*) and a key nutrient that drives susceptibility to fungal infections(*9, 10*). However, the host’s inability to maintain control of iron metabolism increases susceptibility to bacterial(*11, 12*) and fungal infections(*9, 10*). Iron overload worsens outcomes in disseminated *Candida albicans* (*C. albicans*) infections(*13*), but very little is known about the role of iron in the pathogenesis of *C. auris* infections, primarily through in vitro observations. Pyrvinium pamoate, an anthelmintic drug, inhibited the growth of *C. auris*, in part by disrupting iron homeostasis, reducing TCA cycle enzyme activity, and inducing mitochondrial dysfunction(*14*). *C. auris* proteinase and esterase activities, resistance of *C. auris* to fluconazole and caspofungin increase with iron supplementation(*15*), while deferiprone, a clinically approved iron chelator, enhanced the activity of echinocandins and, in a clade-specific manner, amphotericin B(*16*). While these studies highlight the importance of iron acquisition for *C. auris* virulence, they relied on nonphysiological iron sources, lacked mechanistic insight, and had not been validated in vivo.

In the mammalian host, iron is predominantly bound to and shuttled between transferrin (transferrin-bound iron: TBI) and the heme moiety of hemoglobin(*17*). As a long-term co-evolving commensal of humans, *C. albicans* can derive iron from both transferrin(*18*) and heme(*19, 20*). We have previously shown that *C. albicans*(*13*) but not *Aspergillus fumigatus*(*21*) (*A. fumigatus*) worsens outcomes in mice with high transferrin iron saturation. In contrast, *A. fumigatus* utilizes heme as a preferential energy source(*21*), indicating that pathogen-specific preferences for physiological iron species exist in nutritional immunity. However, whether *C. auris* extracts iron from transferrin or causes hemolysis to access heme-bound iron for persistence and virulence remains unknown. Bridging this knowledge gap may help design novel therapeutics to target either the fungal iron acquisition program or host pathways that limit microbial iron availability as potential interventions.

In this study, we investigated whether *C. auris* utilizes physiologic iron sources in vitro and in vivo. We found that *C. auris* causes endothelial injury but does not induce hemolysis of extravascular red blood cells or utilize cell-free heme. These findings were validated in vivo in mice deficient in hemopexin, a heme-scavenging protein. Instead of heme, *C. auris* extracts iron from transferrin-bound iron to support growth and virulence. Compared with vehicle-treated controls, *C. auris*-infected immunocompetent mice with elevated transferrin iron saturation had increased fungal burden and tissue pathology. Consistent with this, *C. auris* mutants lacking *SIT1*, which encodes a siderophore iron transporter, exhibited attenuated growth and reduced pathology in immunocompetent mice, underscoring the importance of iron acquisition for *C. auris* persistence in vivo. We further demonstrate that access to excess transferrin-bound iron induces reactive oxygen species and activates antioxidant programs in *C. auris*, rendering it more resistant to neutrophil-mediated killing. Finally, *C. auris* grown in excess transferrin-iron was resistant to caspofungin, indicating that iron reshapes fungal physiology to drive persistence.

## RESULTS

### Endothelial injury and iron accumulation in *C. auris*-infected kidneys

Kidneys from immunocompetent C57BL/6 mice harbored the highest fungal burden among all organs examined (Fig. 1A-B) as previously described(*22*). These findings are consistent with reports in neutropenic mice(*23–25*), indicating that host immune status does not alter organ tropism during hematogenous dissemination of *C. auris*. Fungal cells were predominantly localized to the renal interstitium (Fig. 1C, Supplemental Fig. 1A). Immunofluorescence staining of kidneys infected with RFP-expressing *C. auris* revealed fungal cells in proximity to, and within the lumen of, proximal tubular epithelial cells (PTECs) (Fig. 1D, Supplemental Fig. 1B). The presence of *C. auris* within these nephron compartments suggests that hematogenously disseminated *C. auris* breaches the renal peri-tubular capillaries.

**Figure 1:**
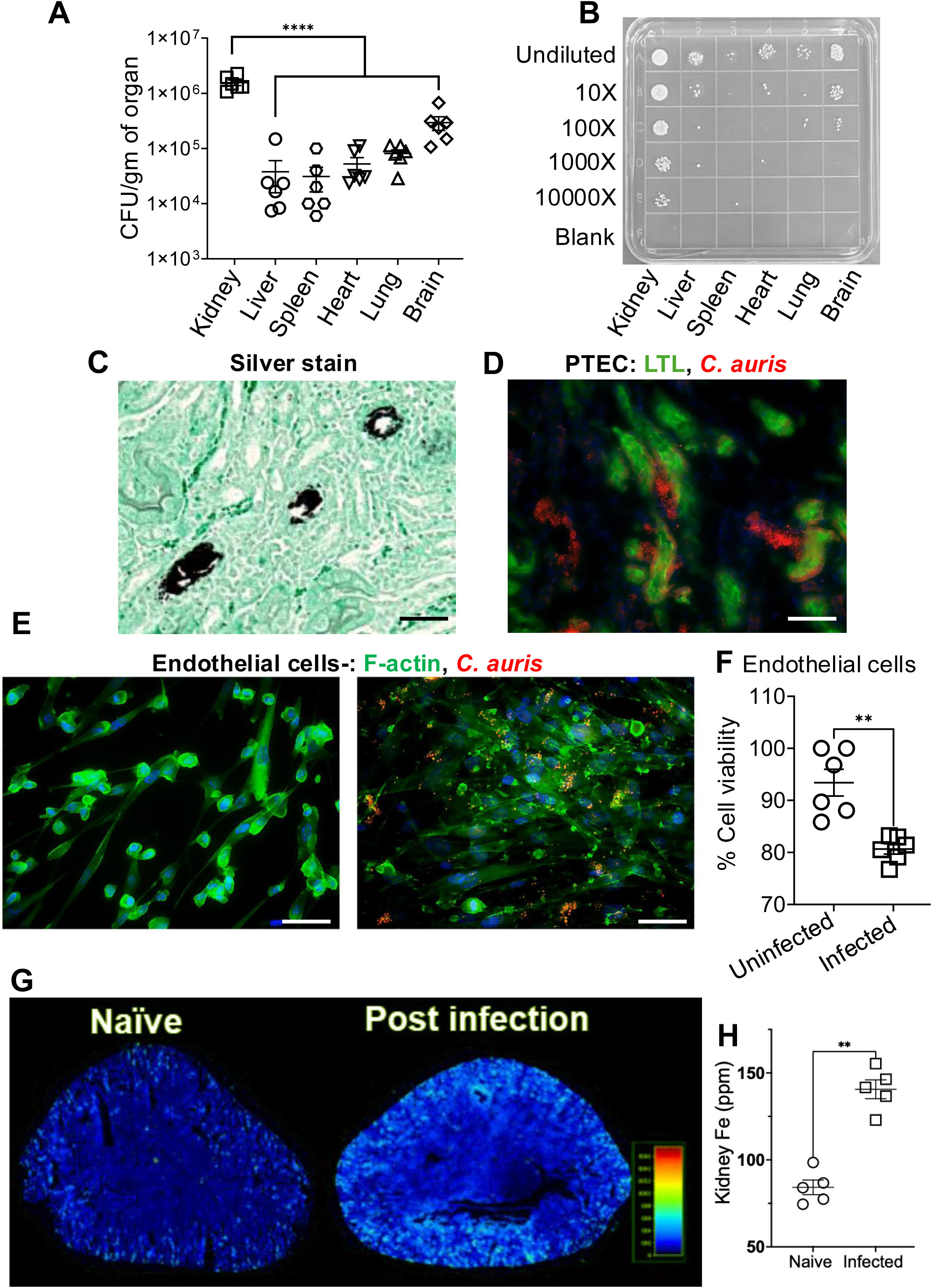
*C. auris* infection is associated with renal intrinsic cell uptake, injury, and iron accumulation. Twelve-week-old immunocompetent C57BL/6 mice were infected intravenously with *C. auris* AR-0382- RFP and outcomes were evaluated after 11 days. Kidneys had the highest fungal burden amongst all the infected organs **(A-B)**. Modified GMS-silver staining revealed C. auris cells in the kidney interstitium (C) and in proximity to, and within the lumen of proximal tubular epithelial cells (PTECs) **(D)**. Mouse primary endothelial cells endocytosed *C. auris* as indicated by colocalization of the *C. auris* RFP and F-actin (green) signals **(E)**. Scale bar 30 μM. This was associated with cell death as indicated by increased LDH released in the co-culture supernatants (Data was normalized to uninfected controls) **(F)**. Experiments were repeated twice, and the pooled data are presented. Unbiased spatial metallomic mass spectrometry of infected mice kidneys revealed increased renal iron accumulation in the cortex and the cortico-medullary regions. A 2-tailed Mann-Whitney test and one-way ANOVA using Tukey’s correction test were used to determine statistical significance. Data are presented as mean ± SEM. **p < 0.005, and ****p < 0.0001

Endothelial traversal may occur via paracellular passage through tight junctions or via transcytosis through endothelial cells—mechanisms described for other pathogenic fungi(*26, 27*) but not previously demonstrated for *C. auris*. To assess whether *C. auris* employs either of these mechanisms, we co-cultured RFP-*C. auris* with primary mouse endothelial cells. After 6 hours, we evaluated fungal transport across an endothelial monolayer grown on 4-µm pore inserts, intracellular fungal localization, and lactate dehydrogenase (LDH) release. No RFP-positive *C. auris* was detected in the medium beneath the insert; however, an intracellular RFP signal was observed within endothelial cells (Fig. 1E). In addition, LDH release was significantly increased in fungus-exposed endothelial cultures, indicating enhanced cell death (Fig. 1F).

Spatial mass spectrometry revealed increased iron accumulation in *C. auris*–infected kidneys, primarily localized to the cortex and corticomedullary regions—areas enriched in PTECs (Fig. 1G-H). Collectively, these data support a model in which, following hematogenous dissemination, *C. auris* may breach the renal vasculature, enter the interstitium, and subsequently be taken up by PTECs, the most abundant epithelial cell type in the kidney(*28*). Endothelial cell death may lead to localized pockets of hemorrhage and hemolysis, providing a mechanistic explanation for iron accumulation in infected kidneys.

### Iron acquisition drives in vitro growth and in vivo persistence

To determine whether iron acquisition is essential for fungal growth and persistence, we generated *C. auris* (AR0382) mutants lacking *SIT1* (a gene coding for *C. albicans* ortholog- siderophore iron transporter) and compared their growth to the parental *C. auris* AR-0382 (WT) in vitro. The *sit1Δ* mutants grew less compared to WT *C. auris* in iron-free media (Fig. 2A-B). Supplementing the media with iron increased the growth rate of WT *C. auris,* whereas the growth of *sit1Δ* mutants in iron-supplemented media was comparable to that of WT *C. auris* grown in iron-free medium (Fig. 2A-B). We next evaluated whether *SIT1* influences the outcomes of hematogenous dissemination of *C. auris* infections in vivo, which, unlike in vitro systems, provides additional nutrients to compensate for iron. Compared to immunocompetent C57BL/6 mice infected with WT *C. auris,* those infected with *sit1Δ C. auris* had reduced renal fungal burden (Fig. 2C) and better renal function as indicated by plasma creatinine levels (Fig. 2D), despite having comparable neutrophil and monocyte infiltrates (Fig. 2E-F). These data highlight the importance of iron uptake in *C. auris* growth, persistence, and associated tissue pathology.

**Figure 2:**
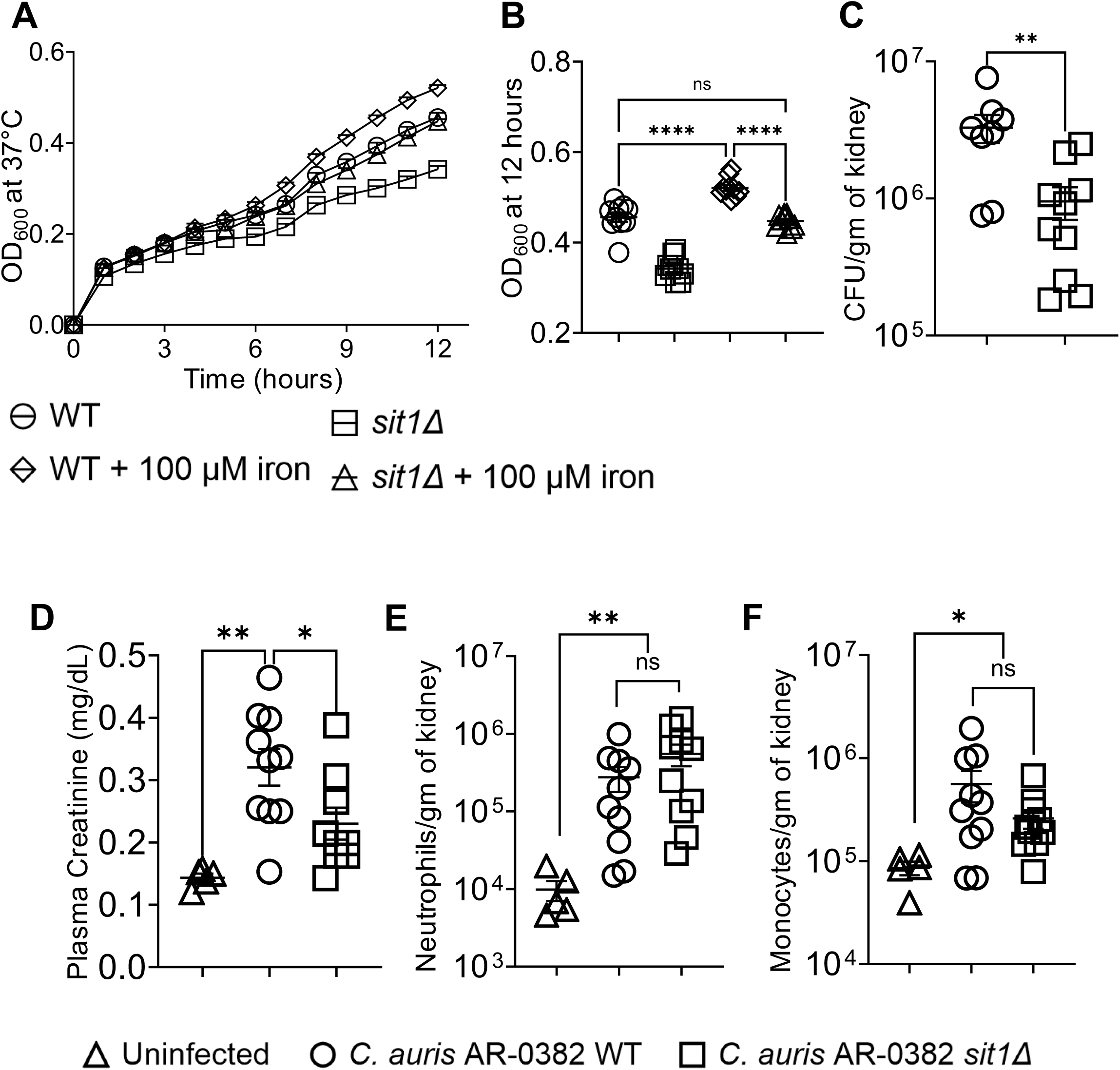
Iron acquisition promotes in-vitro growth and in vivo *C. auris* burden. In vitro, *C. auris* AR-0382 (WT) and *sit1Δ* (impaired siderophoric iron acquisition) were grown in RPMI-1640 with or without 100 μM Fe (ferric ammonium citrate) at 37°C. OD_600_ values measured for 12 hours. *sit1*Δ showed a reduced growth rate in RPMI-1640 compared to WT (**A-B**). 10-to12-week-old immunocompetent C57BL/6 mice were infected intravenously with *C. auris* strain AR-0382 and *sit1Δ* (on the same strain) (5 × 10^7^ yeast per mouse). On day 11, the kidneys of *sit1Δ* infected mice had less fungal burden **(C)**. A lower fungal burden is associated with better preserved renal function, as indicated by lower plasma creatinine levels (**D**). Although *sit1Δ* infection resulted in less injury, the numbers of renal neutrophil and monocyte infiltrates were comparable with the AR-0382 (WT) strain-infected mice **(E-F)**. Experiments were repeated twice, and the pooled data are presented. A 2-tailed Mann-Whitney test and two-way ANOVA using Holm-Šídák’s multiple comparisons test were used to determine statistical significance. Data are presented as mean ± SEM. *p < 0.05, **p < 0.005, and ****p < 0.0001

### *C. auris* does not utilize heme-bound iron

In mammalian hosts, the majority of circulating iron is sequestered within the heme moiety of hemoglobin in red blood cells (RBCs) or bound to the iron-carrier protein transferrin(*17, 29*). Although previous studies have demonstrated a role for iron in *C. auris* antifungal resistance(*15*) and virulence(*30*), these studies relied primarily on iron salts in vitro, and hence the extent to which their findings apply to physiologically relevant host iron sources in vivo remains unclear. To define the contribution of heme- and transferrin-bound iron to *C. auris* pathogenesis, we evaluated fungal growth and virulence-associated traits using physiologically relevant iron sources.

When *C. auris* was cultured with sheep RBCs (1:1 ratio) at 37 °C, it induced minimal to no hemolysis, with levels comparable to the non-hemolytic *C. albicans ece1Δ/Δ* mutant lacking candidalysin. In contrast, the hemolytic *C. albicans* strain SC5314 caused near-complete RBC lysis (Fig. 3A, Supplemental Fig. 1C). To further assess hemoglobin utilization, we measured zones of clearance on sheep blood agar. After 24 hours of incubation, only *C. albicans* SC5314 (positive control) exhibited β-hemolysis, as indicated by halos surrounding each colony. In contrast, all five *C. auris* strains tested (each representing a different clade), as well as the *C. albicans ece1Δ/Δ strain* (negative control), showed no detectable β-hemolysis (Fig. 3B–C, Supplemental Fig. 1D).

**Figure 3:**
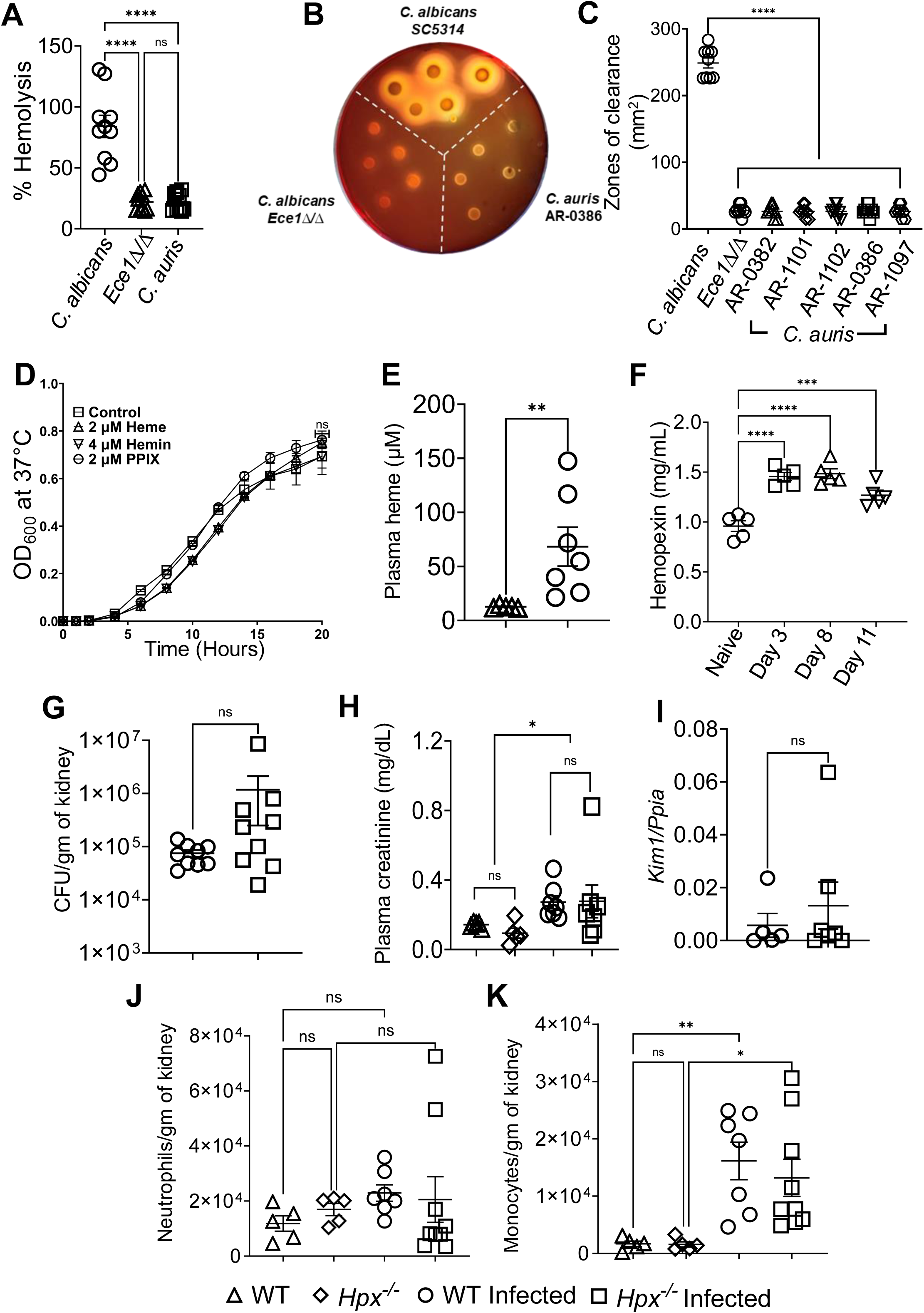
Cell-free heme does not confer a nutritional advantage to *C. auris.* *C. auris* AR-0386, *Candida albicans SC5314*, and *C. albicans Ece1Δ/Δ* were co-cultured with purified sheep red blood cells (RBCs) in a 1:1 ratio for 24 hours. Only C. albicans SC5314 caused RBC hemolysis **(A)** and produced clear zones of β-hemolysis **(B)**. The area of each zone is plotted **(C)**. AR-0382 (Clade I), AR-1101 (Clade II), AR-1102 (Clade III), AR-0386 (Clade IV), AR-1097 (Clade V). *C. auris* AR-0386 was grown in RPMI media supplemented with 2 µM, 4 µM heme, and 2 µM protoporphyrin (organic moiety of heme without iron core) for 20 hours at 37°C. *C. auris* grew comparably in heme or protoporphyrin **(D)**. 10 to 12-week-old immunocompetent C57BL/6 mice were infected with *C. auris* AR0386 intravenously with 4 X 10^7^ yeast/ mouse. Hematogenous dissemination of *C. auris* is associated with increased plasma heme levels **(E)**. Serial tail bleeds showed an increased hemopexin level on day 3, which remained constant till day 11 **(F)**. 10 to 12-week-old immunocompetent Hemopexin knockout (*Hpx^-/-^*) and WT littermates were infected as above. Renal fungal burden (Day 11) was comparable between both strains **(G)**. This was associated with a comparable loss of renal function **(H),** increase in renal injury marker *Kim1* **(I) (***Kim1 gene* expression was not detected in uninjured mice) and neutrophil (**J**) and monocyte (**K**) infiltrates. A 2-tailed Mann-Whitney test and two-way ANOVA using Holm-Šídák’s multiple comparisons test were used to determine statistical significance. Data are presented as mean ± SEM. *p < 0.05, **p < 0.005, ***p < 0.0005 and ****p < 0.0001. All in vitro experiments were repeated twice, with 4-6 replicates per experiment. Mouse experiments were performed twice, and the pooled data are presented.

*C. auris*–induced endothelial injury (Fig. 1E–F) can result in localized pockets of hemorrhage within the infected kidneys. Extravasated RBCs rapidly lyse, releasing cell-free heme (CFH), a known iron source that contributes to the pathogenesis of *C. albicans*(*31*) and *A.fumigatus*(*21*). Whether *C. auris* exploits CFH for growth or virulence remains unclear. To examine this, we compared the growth of *C. auris* (AR0386) in media supplemented with heme, protoporphyrin IX—the iron-free organic backbone of heme(*32*)—or no supplementation. *C. auris* exhibited comparable growth under all conditions, indicating that CFH does not provide a nutritional advantage (Fig. 3D). Collectively, these findings demonstrate that *C. auris* neither secretes hemolytic factors nor degrades and utilizes heme-bound iron, highlighting a fundamental distinction in iron acquisition strategies between *C. auris* and other invasive fungal pathogens.

### Hemopexin deficiency is not a susceptibility factor in disseminated *C. auris* infection

Although CFH did not confer a nutritional advantage to *C. auris* in vitro, growth of this non-commensal microbe in a hostile mammalian host can induce redundant or compensatory iron acquisition pathways that may only be captured in vivo. Plasma heme levels increased during disseminated *C. auris* infection (Fig. 3E). The presence of cell-free heme (CFH) stimulates hepatocytes to upregulate production of hemopexin (Hpx), a high-affinity heme-scavenging glycoprotein(*33, 34*). Consistent with heme release during infection, circulating hemopexin levels were significantly elevated following *C. auris* infection (Fig. 3F).

To evaluate whether the lack of hemopexin and increased in vivo availability of CFH contribute to the pathogenesis of hematogenously disseminated *C. auris* infection, we infected immunocompetent hemopexin-deficient (*Hpx^⁻/⁻^*) mice and WT littermate controls with *C. auris* (AR-0386). Hemopexin deficiency did not alter the outcome of disseminated *C. auris* infection. WT and *Hpx^⁻/⁻^* mice had comparable renal fungal burden (Fig. 3G), renal function (as assessed by plasma creatinine), and tubular injury (*Kim1* gene expression) (Fig. 3H–I). Moreover, renal neutrophil and monocyte accumulation was comparable in WT and *Hpx^⁻/⁻^* mice (Fig. 3J–K). Together, these data indicate that heme-bound iron is dispensable for *C. auris* pathogenesis. Rather, disease progression is driven by non-heme iron acquisition pathways, highlighting a fundamental divergence between *C. auris* and other invasive fungal pathogens that rely on heme-based iron scavenging.

### *C. auris* utilizes transferrin-bound iron as a source of iron during disseminated candidiasis

More than 90% of plasma iron is bound to transferrin, which constitutes the largest circulating reservoir of physiologically available iron(*29*). To determine whether *C. auris* can extract and utilize transferrin-bound iron, we cultured *C. auris* in media supplemented with increasing levels of transferrin iron saturation. Relative to iron-free conditions (basal RPMI-1640), fungal growth was significantly enhanced at 40% transferrin-iron saturation (RPMI-1640 containing 40% Holotransferrin + 60% Apotransferrin), a level typically observed in healthy humans, and further elevated at 90% transferrin-iron saturation (RPMI-1640 containing 90% Holotransferrin + 10% Apotransferrin), which models iron overload states (Fig. 4A). One mechanism that may underlie this enhanced growth, particularly at later time points (Fig 4A), could be an increased dependence on iron to sustain proliferation under nutrient-limited conditions. Ergosterol abundance in the fungal cell membrane correlates with growth(*35*). Fungal cell proliferation and membrane biogenesis require ergosterol synthesis from lanosterol, a process linked to growth rate(*36*) and impaired by iron chelation in *Saccharomyces cerevisiae*(*37*). We observed that *C. auris* grown in 90% transferrin-saturated media exhibited a significantly higher ergosterol-to-lanosterol ratio compared with cells grown in apo-transferrin–containing media (Apo-transferrin: iron-free transferrin was added to maintain constant transferrin levels) (Fig. 4B). Compared to iron-free RPMI, *C. auris* grown overnight in RPMI containing 90% holotransferrin (+10% Apotransferrin) exacerbated primary mouse endothelial cell death (Fig. 4C). To assess the relevance of these findings in vivo, we increased transferrin iron saturation in immunocompetent C57BL/6 mice by administering iron dextran (200 mg/kg), a clinically used iron formulation(*38*), prior to infection with *C. auris* AR-0386. Iron dextran treatment markedly increased transferrin-iron saturation (Fig. 4D) and resulted in iron deposition within both the glomerular and tubular compartments of the kidney (Fig. 4E-F, Supplemental Fig. 2A-B). Elevated transferrin-iron saturation was associated with significantly increased renal fungal burden (Fig. 4G). Notably, fungal growth in the kidney localized predominantly to iron-rich regions (Fig. 4H-I, Supplemental Fig. 2C–D) and was accompanied by substantial renal parenchymal injury, including tubular dilation, loss of nuclear staining, and multiple inflammatory foci (Fig. 4J-K, Supplemental Fig. 2E–F). Compared with vehicle-treated controls, iron dextran–treated mice exhibited impaired renal function, increased tubular injury, and pronounced neutrophil and monocyte accumulation (Fig. 4L–O). Collectively, these data demonstrate that *C. auris* can extract and utilize transferrin-bound iron to support growth and identify elevated transferrin saturation as a host risk factor that exacerbates disseminated *C. auris* infection.

**Figure 4:**
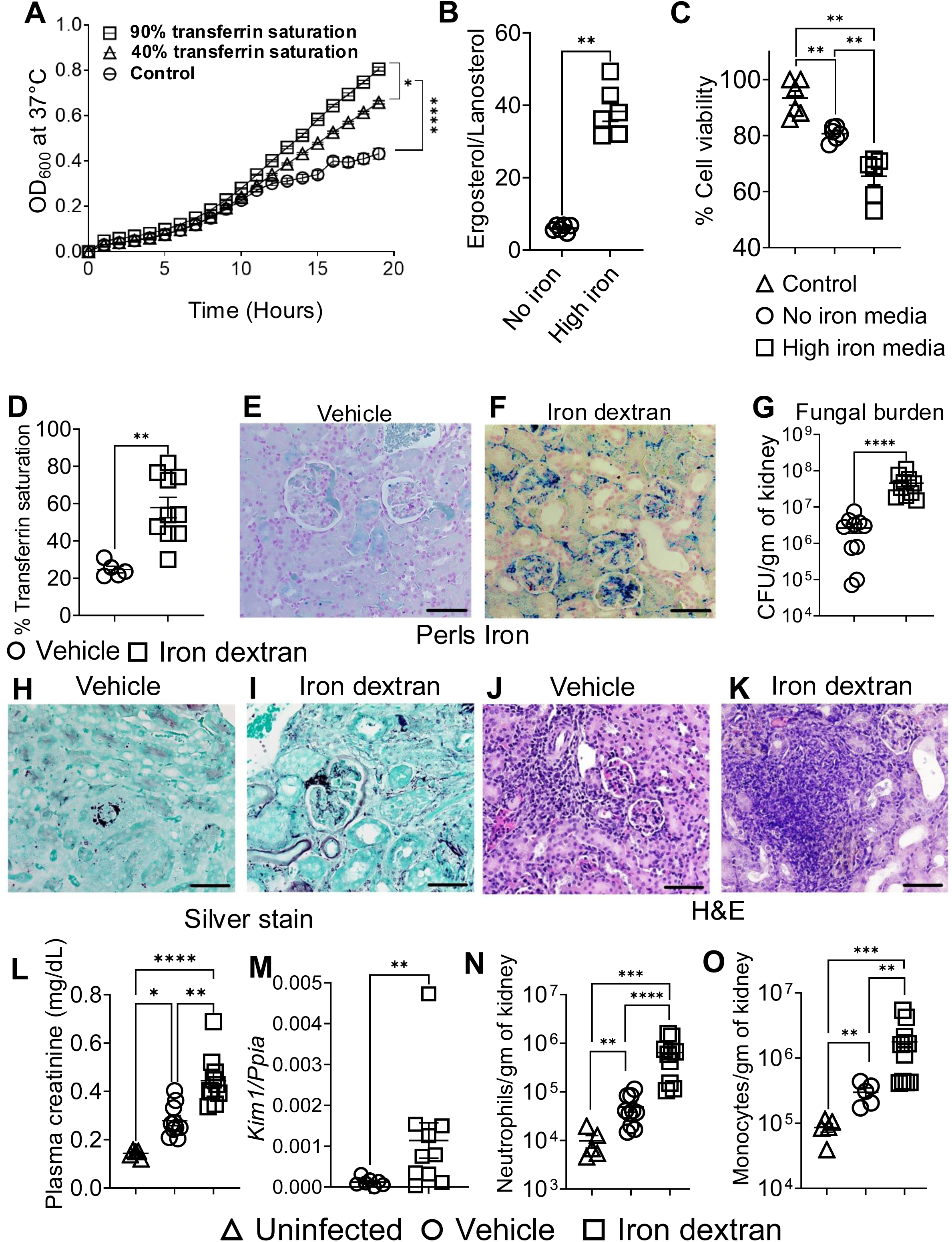
C. auris extracts and utilizes holotransferrin (iron-containing transferrin) as a source of iron for growth and virulence. *C. auris* AR0386 was grown in basal RPMI-1640 and RPMI-1640 supplemented with 40% or 90% transferrin saturated with iron (holotransferrin), for 20 hours at 37°C. Apotransferrin (iron-free transferrin was added to maintain transferrin levels). *C. auris* growth rate increased with transferrin-iron saturation **(A).** Sterol analysis showed an elevated lanosterol/ergosterol ratio in 90% transferrin saturated media as compared to basal RPMI-1640, suggesting iron-driven lanosterol to ergosterol conversion **(B)**. Compared to *C. auris* grown in basal RPMI, *C. auris* grown in RPMI-1640 supplemented with 90% transferrin saturated with iron reduced primary mouse endothelial cell viability, as quantified by an LDH release assay (data are normalized to uninfected controls) (**C**). 10- to 12-week-old male and female immunocompetent C57BL/6J mice were injected with PBS (vehicle) or iron-dextran. Iron-dextran infusion increased transferrin saturation **(D)** and resulted in iron deposition within the glomerulus, tubular, and interstitial compartments of the kidney **(E-F)**. Upon infection with *C. auris* AR0386, mice treated with iron dextran presented with significantly higher renal fungal burden **(G).** Fungal growth was observed in the glomeruli, renal tubules, and the interstitium, regions of iron deposition **(H-I).** Increased fungal burden was accompanied by multiple large foci of inflammation and dilated, a-nuclear tubules with casts **(J-K).** Renal injury quantified as an increase in plasma creatinine levels (**L**), and *Kim1* gene expression (**M**) was increased in the iron-dextran group and was associated with a greater increase in neutrophil and monocyte numbers **(N-O)**. A 2-tailed Mann-Whitney test and two-way ANOVA using Holm-Šídák’s multiple comparisons test were used to determine statistical significance. Data are presented as mean ± SEM. *p < 0.05, **p < 0.005, ***p < 0.0005 and ****p < 0.0001. All experiments were repeated twice, with 5-6 replicates for each experiment. Mouse experiments were performed twice, and the pooled data are presented.

### Transferrin-bound iron promotes *C. auris* virulence

To determine whether transferrin-bound iron influences *C. auris* physiology beyond its role in cell proliferation, we cultured *C. auris* AR0382 in basal RMPI-1640 or in RPMI-1640 containing 90% transferrin-saturated iron and quantified intracellular reactive oxygen species (ROS). Exposure to iron resulted in a rapid increase in intracellular ROS levels (Fig. 5A), indicating that iron availability induces oxidative stress in *C. auris*. This finding led us to posit that iron-generated ROS may activate antioxidant defense pathways, enabling *C. auris* to adapt to oxidative environments. Consistent with this hypothesis, *C. auris* grown in 90% transferrin-iron saturated medium, but not in basal medium, showed significant upregulation of *SOD4* (superoxide dismutase 4; *C. albicans* ortholog) and *AOX2* (alternative oxidase 2; *C. albicans* ortholog) (Fig. 5B–C). To assess the functional relevance of iron-induced antioxidant priming, we evaluated susceptibility to neutrophil-mediated killing. *C. auris* grown under low-iron or 90% transferrin-iron–saturated medium conditions was opsonized in RPMI supplemented with 10% FBS and incubated with freshly isolated neutrophils for 3 hours (MOI 1:3, neutrophils: fungi). Following incubation, neutrophils were lysed, and surviving fungi were quantified by colony-forming unit (CFU) enumeration. *C. auris* cultured in high-iron conditions exhibited significantly increased resistance to neutrophil-mediated killing, as evidenced by higher CFU recovery (Fig. 5D). Together, these indicate that exposure to iron may trigger a ROS-mitigation program in *C. auris*, thereby increasing resistance to neutrophil-mediated killing.

**Figure 5:**
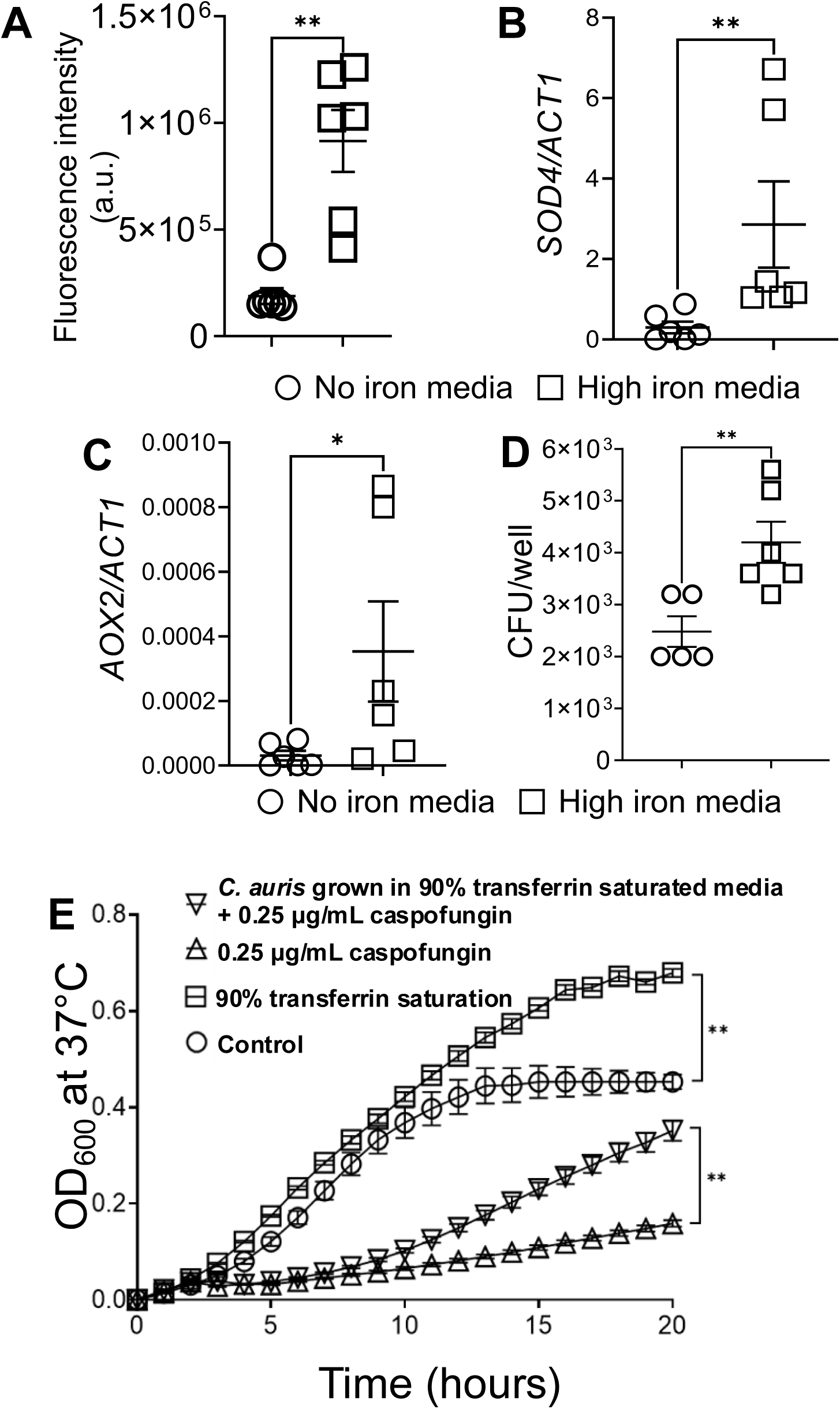
Uptake of transferrin-bound iron increases *C. auris* resistance to neutrophil and caspofungin-mediated killing. *C. auris* AR-0382 was grown in basal RPMI-1640 or 90% transferrin-iron saturated RPMI-1640 (high iron medium) at 37°C. After 30 minutes, ROS was induced in *C. auris* grown in high-iron medium **(A)**. Continued growth under the above conditions for 6 additional hrs was associated with increased gene expression of *SOD4* (Superoxide Dismutase 4) and *AOX2* (Alternative Oxidase 2) **(B-C)**. These iron-primed *C. auris* cells were resistant to neutrophil-mediated killing, as indicated by increased CFUs after co-culture with neutrophils **(D)**. Additionally, *C. auris* AR0382 (a caspofungin-susceptible strain), grown in 90% transferrin-iron saturated RPMI-1640, was resistant to caspofungin-mediated killing **(E)**. A 2-tailed Mann-Whitney test and two-way ANOVA using Holm-Šídák’s multiple comparisons test were used to determine statistical significance. Data are presented as mean ± SEM. *p < 0.05, **p < 0.005. All experiments were repeated twice, with 3-6 replicates in each experiment.

Strains originating from *C. auris* clade 1 are susceptible to caspofungin(*39, 40*). Hence, we also evaluated whether AR0382, grown in high transferrin-iron saturation, is resistant to caspofungin. AR0382 was grown under identical conditions described for neutrophil studies. After washing to remove excess iron, the fungus was added to basal RPMI containing 0.5μg/mL caspofungin, and the growth rate was monitored. Compared to *C. auris* AR0382 grown in basal RPMI, the fungus grown in 90% transferrin-iron medium grew significantly more despite the presence of caspofungin and was comparable to *C. auris* grown in basal RPMI in the absence of caspofungin (Fig.5E). Taken together, these findings indicate that elevated transferrin-iron saturation can promote a caspofungin-resistant growth phenotype in AR0382, suggesting that host-iron dysregulation may serve as a critical driver in modulating anti-fungal resistance.

## DISCUSSION

*C. auris* is strikingly resistant/tolerant to antifungal drugs, which results in poor clinical outcomes(*6, 41, 42*), and the limited pipeline of antifungal drugs necessitates identifying host conditions that promote antifungal resistance and susceptibility to infection, thereby opening opportunities for intervention. Recent in vitro studies identified iron as a key regulator of *C. auris* fitness and resistance to azoles (*15*) and echinocandin(*16*). Furthermore, iron binding to caspofungin reduces its ability to inhibit the β-1,3-D-glucan synthase in *C. albicans*(*43*), a mechanism not evaluated in *C. auris*. Collectively, these findings highlight targeting host iron metabolism as a novel, underexplored therapeutic approach that is distinct from conventional antifungal strategies. In this study, we use cell culture systems, immunocompetent mice with iron overload, and *C. auris* mutants with impaired iron uptake to demonstrate that *C. auris* can extract iron from transferrin but not from cell-free heme, thereby supporting growth and virulence. This contrasts with *C. albicans, A. fumigatus, and C. neoformans,* fungi known to utilize heme(*44–46*), and highlights a key metabolic divergence among fungal species.

Following hematogenous infection, *C. auris* is predominantly observed in the renal interstitium, indicating that it may breach the vascular endothelial barrier. Indeed, we demonstrate that *C. auris* invades the endothelial cells by transcytosis and induces endothelial cell death. The inability to traverse the endothelial layer via paracellular routes suggests a lack of *C. albicans* or *C. neoforma*ns-like hydrolytic enzymes, such as secreted aspartic proteinases and lipases that directly cleave tight junction proteins(*26, 27*). The endothelial cell death is exacerbated by *C. auris* exposed to iron, suggesting that excess iron availability may remodel *C. auris*, making it more immunogenic and virulent. In support, we have previously shown that ferric ammonium citrate-containing medium sustains *C. albicans* hyphae with exposed β-1,3-glucan and accentuates IL-8 production in human renal epithelial cells(*13*).

Injury to the endothelium results in multiple microhemorrhages and the local release of extravascular RBCs, which rupture and release cell-free heme(*47*), potentially explaining the increased iron deposits in the *C. auris*-infected kidneys. Unregulated accumulation of CFH is toxic to the renal parenchyma(*48*), signaling hepatocytes to increase hemopexin production(*49*). We have previously shown that hemopexin knockout mice do worse following *A. fumigatus* infection(*21, 50*), an observation not recapitulated in *C. auris* infection. *A. fumigatus* produces the Asp-hemolysin, a hemolytic toxin(*51*) and thrives in cell-free heme(*21*). We now identify that *C. auris* is unable to hemolyze RBCs or utilize heme to gain a nutritional advantage (Fig. 3). These observations suggest that *C. auris* neither secretes a hemolytic factor nor has an active heme uptake or breakdown machinery. As *C. auris* is a new pathogen of humans, it may not have evolved mechanisms to use heme as a major iron source. The increase in hemopexin during *C. auris* infection is probably due to its role as an acute-phase reactant, independent of hemolysis(*52*).

In contrast to heme, *C. auris* actively extracts iron complexed to transferrin, as indicated by its higher growth in a 30% transferrin-iron-saturated medium, which is further accentuated when transferrin iron saturation reaches 90% an observation recapitulated in vivo. Multiple *C. auris* clades upregulate the siderophoric iron uptake transporter *SIT1* gene, which mediates the uptake of iron-containing siderophores(*53*). The attenuated growth of *SIT1* mutants reinforces the importance of limiting iron uptake in *C. auris* infections.

Increased ROS in neutrophils correlates with increased killing of *C. auris*(*54*). Neutrophil depletion worsens outcomes and mortality in *C. auris-*infected mice(*24*), but iron-dextran-injected mice, despite being neutrophil-sufficient, had worse outcomes. We now find that iron supports growth and makes *C. auris* resistant to neutrophil-mediated killing. At a mechanistic level, iron induced ROS in *C. auris* and activated the transcription of genes associated with antioxidant defense. In *C. albicans*, Sod4 and Sod5 detoxify extracellular ROS produced by innate immune cells(*55*), and Sod5 is required for resistance to neutrophil-mediated killing(*56*). Notably, *C. auris* Sod4 exhibits functional similarity to *C. albicans* Sod5(*57*), suggesting that iron-induced *SOD4* expression may enhance the capacity of *C. auris* to neutralize host-derived oxidative stress-mediated killing. Similarly, *AOX2* in *C. albicans* has been shown to alleviate elevated ROS induced by high-iron media(*58*).

It was recently reported that iron binding induces conformational changes in caspofungin and reduces its efficacy against *C. albicans*. Caspofungin was equally effective against *C. albicans* grown overnight in normal- or high-iron medium. However, we observed that *C. auris* AR0382, a caspofungin-susceptible strain (*59*) grown overnight in 90% transferrin-saturated medium, partially resisted caspofungin-mediated killing, supporting the notion that iron alters *C. auris* beyond its obligatory role in proliferation. Rapid iron-induced growth may also trigger functional diversity in cell wall proteins, as evidenced by the ability of *C. auris* grown in a high-iron medium to exacerbate endothelial cell death.

Collectively, these findings support a model in which transferrin-bound iron induces an oxidative stress response in *C. auris* that preconditions the fungus by upregulating antioxidant defenses and thereby enhancing survival in the face of neutrophil-derived ROS. This iron-dependent priming mechanism provides a direct link between host iron availability and increased *C. auris* virulence.

### Limitations of the study

*C. auris* isolated from a mammalian host undergoes modifications in cell-surface antigens and a metabolic switch, which is genetically conserved in daughter cells(*60*). Whether growth in an iron-rich environment triggers such modifications is not investigated in this study. Such changes aid in avoiding immune surveillance and in survival and persistence within the host(*61*). A further caveat when working with transferrin saturation above ∼85% is the emergence of non–transferrin-bound iron(*62, 63*), a highly bioavailable iron source for microbes(*52, 64, 65*). Dissecting the contribution of specific iron species to *C. auris* pathogenesis is essential for defining mechanism-based risk factors and informing evidence-based guidance for high-risk patients and warrants future investigations.

### Summary

Our findings underscore the key differences in the iron-acquisition mechanisms of this emerging pathogen and can help inform prognostic strategies for susceptible patients. Of clinical relevance, patients with chronic liver disease exhibit elevated serum iron and transferrin iron saturation(*66–68*), even without hemochromatosis(*69*). This population is at higher risk for *C. auris* infection than patients without liver disease(*70–72*). While this work lays a foundation, further work is required to understand the host’s intrinsic defects and the fine mechanisms by which high transferrin iron saturation influences the outcomes of *C. auris* infections.

## MATERIALS AND METHODS

### Mice

All experiments were performed following the National Institutes of Health and Institutional Animal Care and Use Guidelines and were approved by the Animal Care and Use Committee of the University of Florida (IACUC# 202400000692). The mice were maintained at 23°C (±2°C), with 30–70% humidity and a 14:10 light: dark cycle. Wild-type (WT) and hemopexin knockout (*Hpx^-/-^*) mice were purchased from Jackson Laboratory and maintained in the animal facilities at the University of Florida on a regular diet. An equal number of 10- to 12-week-old male and female mice were used throughout the studies.

### Fungal strains

The *C. auris* strain *AR-0386 (Clade 3)* and *AR-0382 (Clade 1)* used in this study were obtained from the Centers for Disease Control and Prevention’s Antibiotic Resistance Isolate Bank *C. auris* panel. All strains were streaked on Sabouraud Dextrose agar (Difco, DF0747-17-9) at 30°C for 24 hours. RFP-*C. auris* (AR0-382) was generated and characterized in our previous study(*59*). Single colonies were picked from the plates and grown in Sabouraud Dextrose broth (Difco, DF0382-17-9) at 30°C and 300 RPM for 2 overnight cultures. OD_600_ was measured to verify the culture’s growth phase. Log-phase cells (OD_600_ =0.6) were used for infection. Fungal mutant construction and details in the supplemental text(*59, 73*).

### Mouse model of transferrin saturation

Immunocompetent male and female C57BL/6 mice were injected intraperitoneally with vehicle (PBD) or iron dextran (diluted in sterile PBS, Sigma-Aldrich, D8517) at 200 mg/kg on alternate days for 4 weeks as described by Li et.al., 2017(*74*). Plasma was collected weekly to measure serum iron levels using an iron assay kit (Abcam, ab83366). Transferrin saturation was measured as described by Lee et al(*75*).

### Mouse model of candidiasis

Log-phase yeast cells were centrifuged at 1500 RPM for 5 minutes, washed in sterile PBS. 5 × 10^7^ *C. auris* (AR0386) yeast cells were injected intravenously into *Hpx^-/^*^-^, WT littermates, and WT mice with iron overload. In some experiments, mice were infected with equal number of the RFP-labeled *C. auris* AR0382(*59*). *sit1Δ* mutants were generated in the *C. auris* AR0382 background. Mice were euthanized 11 days post-infection. Fresh tissue was used to determine fungal burden and for flow cytometry. Snap-frozen tissue was used for RNA isolation and protein analysis, and 10% formalin-fixed or 1% Periodate-Lysine-Paraformaldehyde fixed tissue was used for histopathology and immunofluorescence.

### Primary endothelial cell culture

C57BL/6 Mouse Primary Lung Microvascular Endothelial Cells were obtained from Cell Biologics (Cell Biologics, C57-6011). Cells were cultured in rat tail collagen-coated flasks and maintained in Endothelial Cell medium (Cell Biologics, M1168) supplemented with 0.5 mL VEGF, 0.5 mL Heparin, 0.5 mL EGF, 0.5 mL FGF, 0.5 mL Hydrocortisone, 5 mL Antibiotic-Antimycotic solution, and 25 mL Heparin at 37 °C and 5% CO_2_. Cells were used between passages 3 and 5.

### *C. auris* endothelial cell interaction and LDH measurement

Primary mouse endothelial cells were grown on 4-micron tissue culture inserts and *C. auris* (AR0386) was added at a multiplicity of infection (MOI) of 1 cell : 6 yeast for 6 hours. After the incubation period, the secreted LDH was measured using an LDH assay kit (Cayman chemical, 601170) according to the manufacturer’s instructions.

### Fungal growth curve

*C. auris* cells were grown overnight in Sabouraud dextrose broth in a shaking incubator (200 RPM) at 30 °C. Twelve to fourteen hours later, the fungal cells were centrifuged, washed in sterile PBS, and reconstituted in RPMI-1640 (Corning, 10-104-CV) without any supplementation (no-iron media). 10,000 yeast cells were added to each well. For the heme growth curve, RPMI-1640 was supplemented with 2 μM, 4 μM heme, and 2 uM protoporphyrin IX (Frontier Specialty Chemicals, P562-9). High iron media was prepared by supplementing RPMI-1640 with human holo-transferrin (Sigma Aldrich, T0665) containing 25 μM iron (∼30% transferrin saturation) or 100 μM iron (∼85% transferrin saturation). Human apo-transferrin (Athens research, 16-16-A32001-BPG) was added to maintain a constant total transferrin concentration of 50 μM transferrin. For caspofungin resistance, *C. auris* cells were grown with or without transferrin overnight. Cells were split into media supplemented with or without 0.5 μg/mL caspofungin diacetate (Sigma Aldrich, SML0425). The growth rate was monitored for 20 hours at 37°C with orbital shaking. OD_600_ was recorded at 60-minute intervals and plotted as mean ± SEM. Each experiment was repeated twice, with 6 replicates per condition.

### Blood agar culture

Blood agar plates were prepared using a mixture of 5% sheep red blood cells and 1.5% agar supplemented with 3% D-glucose (Sigma-Aldrich, G7021). 100,000 yeast cells (in 10 μL) were spotted on the plate and incubated for 24 hours at 37°C. The zone of clearance is expressed as the area of the circle in mm^2^.

### Hemolysis assay

Hemolytic activity of *C. auris* was measured by co-incubation it with RBCs as described by Mogevero et al(*45*). Briefly, yeast and RBCs were incubated together at a 1:1 ratio at 37℃ and 50 rpm for 24 hours. After 24 hours, the tubes were centrifuged, and the supernatant was transferred to a 96-well plate for hemoglobin measurement at 414 nm. Data was normalized to the absorbance of RBCs lysed with water and plotted as percent hemolysis.

### *C. auris* ergosterol and lanosterol measurement

*C. auris* primary culture was grown as described above. The secondary culture was split into two flasks, one containing no iron and the other 100 μM holo-transferrin and incubated for 24 hours. Post-incubation, samples were centrifuged, air-dried, and weighed. Ergosterol and lanosterol levels were measured using UPLC-MS/MS. The protocol is described in the supplemental text(*76*).

### *C. auris* and neutrophil co-culture ex vivo

*C. auris* primary culture was grown as described above and split into a secondary culture that was grown in the absence of iron or in 100 μM holo-transferrin, as above. After washing, the yeast cells were opsonized in RPMI-1640 medium supplemented with 10% FBS for 45 minutes. Neutrophils were isolated from 10-week-old C57BL/6 mice using the EasySep Mouse neutrophil isolation kit (STEMCELL Technologies, 19762). Opsonized *C. auris* was cultured with purified neutrophils at an MOI of 1:3 (yeast: neutrophil) for 3 hours. After incubation, neutrophils were lysed with 0.02% Triton in water. Fungal cells recovered from neutrophils were plated on Sabouraud dextrose agar plates for CFUs per well, as described above.

### Fungal burden determination

Mice were euthanized with 1.5% (w/v) avertin and bled via cardiac puncture. The kidneys were harvested aseptically, weighed, and homogenized with a tissue homogenizer. Tissue homogenates were serially diluted and plated onto Sabouraud dextrose agar. Colony-forming units (CFUs) were counted after 24 hours of incubation at 37 °C, and results were expressed as CFUs/gram of kidney.

### Flow cytometry

Single-cell suspensions from the kidneys were prepared as described previously(*13*). Single cells were incubated for 20 minutes with eBioscience viability dye eF780 (1:3000 in PBS) and anti-CD16/32 (FC block, 1:100). After washing with FACS buffer, cells were stained for CD45 (PerCP conjugate, Clone 30-F11, Biolegend, 103130), CD11b (APC-Fire 810 Clone M1/70, Biolegend, 101287), Ly6G (Spark UV 387, Clone 1A8, Biolegend, 127677), and Ly6C (Brilliant violet 711, Clone HK 1.4, Biolegend, 128037). Flow cytometry data were obtained using a Cytek Aurora 5-laser system (Fremont, CA) and analyzed using FlowJo software 9.0.

### Perl’s detectable iron staining

Five-micron-thick paraffin-embedded tissue sections were used to detect Perl’s iron deposits as we described(*77*). The tissue was counter-stained with nuclear fast red and imaged.

### H&E staining

10% formalin-fixed and paraffin-embedded 5-μm-thick sections were stained with hematoxylin and eosin (H&E) as previously described(*13, 77*). Briefly, the sections were deparaffinized with xylene, stained with CAT’s hematoxylin for 3 minutes, and with eosin for 30 seconds. The dehydrated slides were mounted with Cytoseal XYL and imaged.

### Silver stain

Paraffin-embedded sections were used to detect fungal location and morphology as described previously(*13*), using a commercial silver-staining kit (Thermo Fisher, HT100A) according to the manufacturer’s instructions.

### Immunofluorescence

Mouse primary endothelial cells were cocultured with *C. auris* AR0382-RFP on glass coverslips as described above. After the incubation period, cells were fixed with 4% PFA and stained using Acti-stain 488 phalloidin (Cytoskeleton Inc. PHDG1-A) as described by the manufacturer. Immunocompetent C57BL/6 mice were infected with *C. auris* RFP (AR-0386) intravenously. Mice were euthanized after 4 days. 1% Periodate-Lysine-Paraformaldehyde fixed kidney sections (five micron thick) were used for the immunofluorescence detection of RFP-tagged *C. auris* and lotus tetragonolobus lectin (LTL, Vector Laboratories, FL-1321-2) positive proximal tubular epithelial cells, as described previously(*13*).

### Real-time PCR

RNA from snap-frozen kidney sections was purified as previously described(*13, 77*). One μg of RNA was used to synthesize cDNA using iScript cDNA synthesis kit (Bio-Rad Laboratories, 1708841). The cDNA template was mixed with SSO advanced SYBR Green Universal Supermix (Bio-Rad Laboratories, 1725271) for quantitative PCR. Relative expression for *Kim1* (BioRad, qMmuCIP0030545) was calculated using 2^(-ΔcT)^ method. *Ppia* (BioRad, qMmuCED0041303) was used as the housekeeping/reference gene for mouse genes.

For *C. auris* RNA extraction, cells were resuspended in TRIzol and bead-beaten for 10 minutes. The bead tubes were centrifuged, and the supernatant was removed into new tubes. The method described above was followed for RNA isolation, cDNA synthesis, and qPCR. Relative expression for *SOD4* (F: TTGGAAACCCTTCACTTTCG, R: TCAGATGAGCAAGGCGCCA) and *AOX2* (F: GCTATCCATCAATAAGCC, R: CAAATCTTCTTTCTCCCAGC) was calculated using 2^(-ΔcT)^ method. *ACT1* (F: CGTGCTGTGTTCCCATCCAT, R: AGCCTCATCACCGACATACG) was used as the housekeeping/reference gene for *C. auris*.

**The details for imaging mass spectrometry are provided in the supplementary methods section**(***78–80***).

## Author contributions

Y.S. conceived the study and designed the experiments. T.A, D.K, T.A, K.H., G.G., S.H., and Y.S performed the experiments. G.Z and T.R.O generated and provided C. auris AR0382-RFP and AR0382 sit1Δ strains. A.G and A.S provided the quantification of ergosterol and lanosterol. T.P analyzed the kidney section for spatial mass spectrometry. M.S.L provided experimental protocols. T.R.O and M.S.L discussed the results and gave expert advice. T.A., D.K., and Y.S analyzed the data. T.A and Y.S wrote the original draft of the manuscript. All authors edited the manuscript and approved the final version.

---

**Supplementary figure 1.**
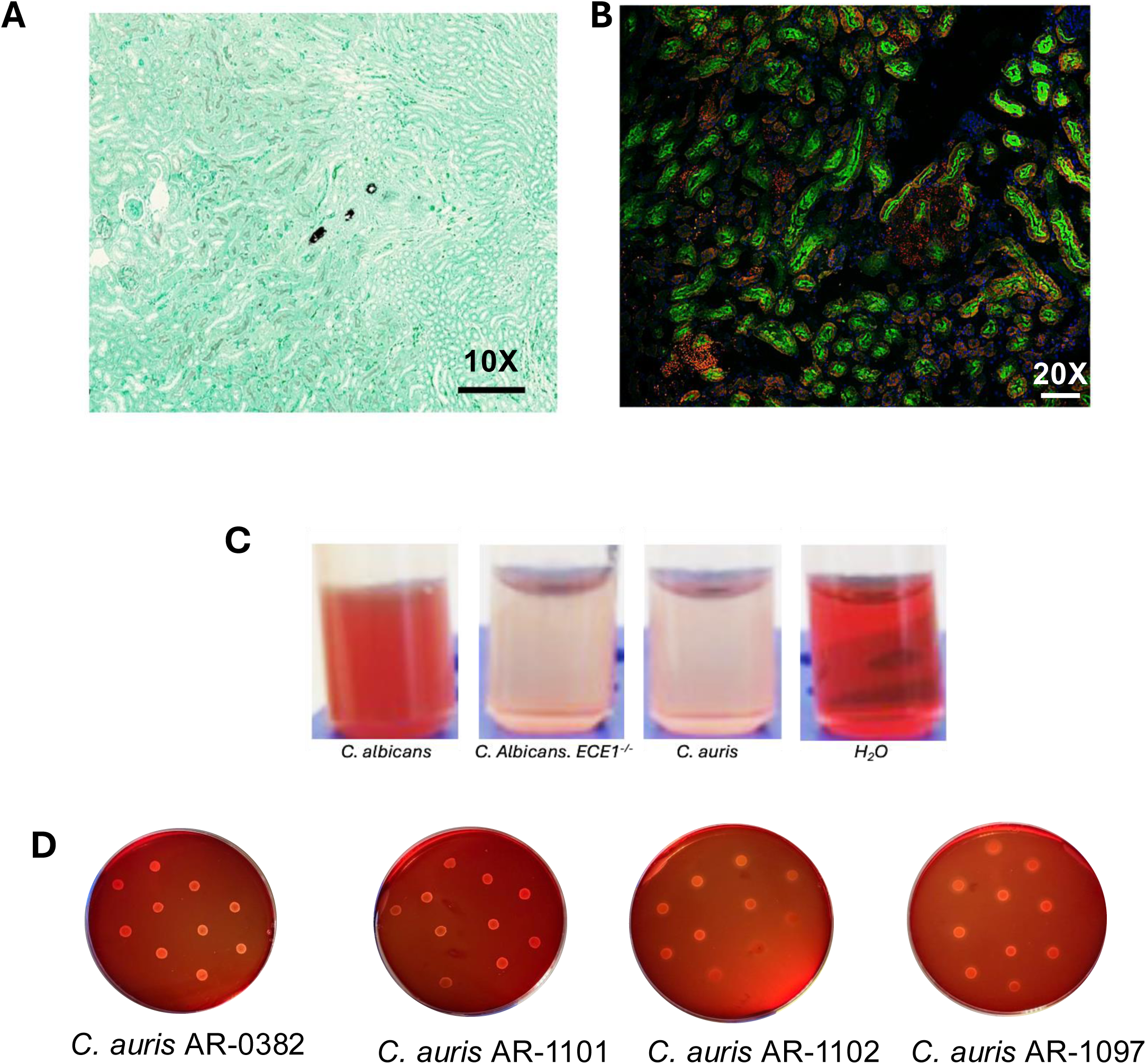
An equal number of 10-12-week-old male and female C57BL/6J mice were infected with RFP-labeled 5 X 10^7^ *C. auris* (AR0382, intravenous), and the tissue was collected 11 days later. Silver staining of formalin-fixed sections showed the presence of *C. auris* yeast cells in the kidney interstitium **(A)**. Kidney sections frozen in 4% PLP showed the presence of RFP-labeled *C. auris* in close proximity of lotus tetragonolobus lectin positive proximal tubular epithelial cells **(B)**. Scale bar: at 10X- 100 μM; 20X- 50 μM. *C. auris* AR-0386, AR-1101, AR-1102, AR-1097, *C. albicans SC5314, and C. albicans Ece1Δ/Δ* were grown in Sabouraud dextrose broth overnight at 30 °C. An equal number of yeast cells and RBCs were co-cultured at a 1:1 ratio, at 37°C and 50 rpm for 24 hours. After 24 hours, hemoglobin was measured in the supernatant at 414 nm (**C**). 100,000 yeast cells from the same culture were plated on blood agar plates supplemented with 3% glucose and incubated at 37°C for 24 hours to measure β-hemolysis (**D**).

**Supplemental Figure 2.**
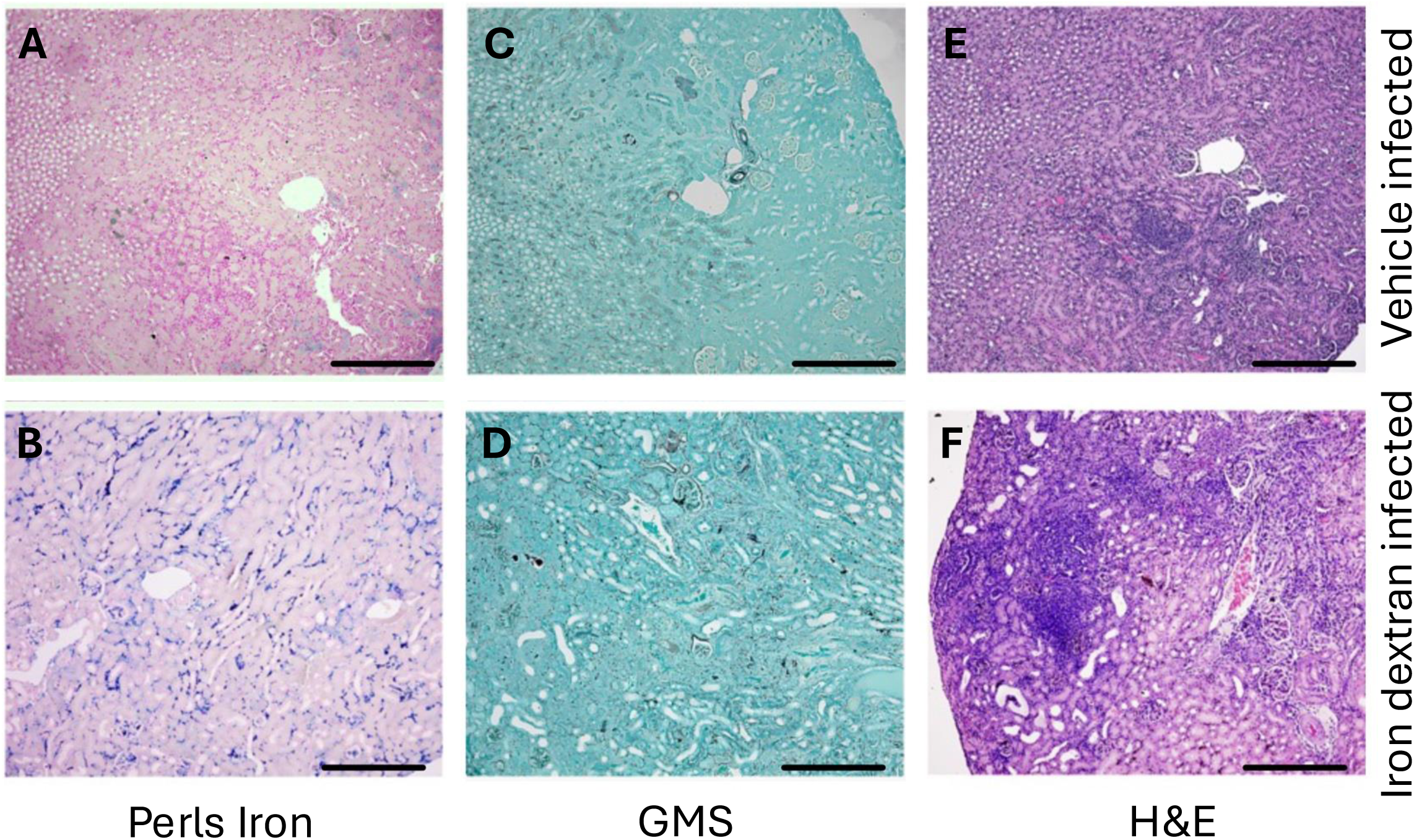
Equal number of 10-12-week-old male and female C57BLk/6J were injected with PBS (vehicle) or Iron-dextran diluted in vehicle (200mg/kg, 2 times a week for 3 weeks). Mice were infected with 5 X 10^7^ *C. auris* (AR0386, intravenous), and the tissue was collected after eleven days. Compared to the vehicle, the kidneys of iron-dextran-treated mice had large deposits of Perls-detectable iron (**A-B**). This was associated with increased fungal growth (Grocott modified silver (GMS) staining) (**C-D**). H&E staining revealed large foci of inflammation and tubular casts in the iron dextran-injected mice (**E-F**). Scale bar : 100μM.

