## Supplemental Methods for "Candidozyma auris utilizes transferrin, but not heme-bound iron for in vivo virulence"

Tanmay Arekar *et al.*

**This PDF file includes:**

Supplementary Text
Method details
Tables S1 to S4
References (59,73,76,78-80)

**Supplementary Text**

***C. auris* mutant construction**

*C. auris* strains were generated with a transient CRISPR system *(59)*. The universal CAS9 cassette was amplified from plasmid pTO135 using oTO143-oTO41. The gRNA cassette used to make *SIT1* mutant strain was constructed using a Splice-On-Extension reaction, incorporating a 20-bp specific gRNA targeting the *SIT1* gene in primers oTO2967- oTO2968. The gRNA cassette used to make *FTR1* mutant strain was constructed using the Splice-On-Extension reaction with primers oTO2979- oTO2980. The repair templates for making the mutants were maintained in the multiple cloning site of the pUC19 cloning vector *(73)* and assembled from fragments as described below using the NEBuilder HIFI DNA Assembly master mix (NEB #E2621) according to the manufacturer's instructions.

To assemble pTO458 (*pSIT1*::*NAT*), 500 bp immediately 5' of *SIT1* (*B9J08_002110*) and 500 bp immediately 3' of *SIT1* were amplified from AR0382 genomic DNA using primers oTO2959-oTO2960 and oTO2961-oTO2962 respectively; the *NAT1* expression cassette was amplified from pTO100 using primers oTO669-oTO668; the pUC19 vector backbone was amplified from plasmid pTO139 using primers oTO278- oTO279. The repair cassette was amplified from pTO458 using primers oTO18-oTO19.

**Table S1: Fungal strains:**

| Strain | Genotype | Source |
|---|---|---|
| TO33 | AR0382 | CDC AR bank |
| | AR0382- RFP | (Santana and O'Meara 2021) *(59)* |
| TO944 | AR0382 Δ*sit1 (B9J08_002110)::NAT* | This study |
| TO949 | AR0382 Δ*FTR1 (B9J08_002108)::NAT* | This study |

**Table S2: Plasmids:**

| Name | Description | Source |
|---|---|---|
| pTO135 | *pENO1-CaCas9-tCyc1, AMP* | (Santana and O'Meara 2021) *(59)* |
| pTO136 | *pADH1-tRNA-Ala-gRNA-tracrRNA-HDV-tAgTEF2, AMP* | (Santana and O'Meara 2021) *(59)* |
| pTO139 | pUC19, *AMP* | (Santana and O'Meara 2021) *(59)* |

| | | |
|---|---|---|
| pTO458 | *sit1 (B9J08_002110)::NAT, AMP* | This study |

**Table S3: Oligos:**

| Name | Sequence | Purpose |
|---|---|---|
| oTO41 | GTCCCAAAACCTTCTCAAGC | To amplify Cas9 |
| oTO143 | CCTCTTTGTAGTTCAACTTATGC | |
| oTO18 | CAGGAAACAGCTATGAC | To ampligy repair cassette |
| oTO19 | GTAAAACGACGGCCAG | |
| oTO278 | GGGGATCCTCTAGAGTCG | To amplify pUC19 backbone |
| oTO279 | GGGTACCGAGCTCGAATTC | |
| oTO668 | ACTGGATGGCGGCGTTAG | To amplify *NAT* expression cassette |
| oTO669 | CGACATGGAGGCCCAGAATAC | |
| oTO2967 | GGTATTTAGAGCATCAGTGGGTTTTAGAGCTAGAAATAGCAA | To assemble gRNA targeting *SIT1* |
| oTO2968 | CCACTGATGCTCTAAATACCTGGACGAGTCCGGATTCGAACC | |
| oTO2959 | GTGAATTCGAGCTCGGTACCCTCCAACGGCAATACCTAC | To amplify 500bp 5' homology of *SIT1* |
| oTO2960 | GGGGACGAGGCAAGCTTGATACTCTCAATATTGACTGAAAAATTAAG | |
| oTO2961 | TACTAACGCCGCCATCCAGTGCTTGCTGAAGCTAACAC | To amplify 500bp 3' homology of *SIT1* |
| oTO2962 | AGGTCGACTCTAGAGGATCCTAGTCAGAGAAGGCAAATC | |
| oTO2974 | AGGTCGACTCTAGAGGATCCCCTCCCTCTTGGATGATG | |

**Ultra-performance liquid chromatography-triple quadrupole mass spectrometer (UPLC-MS/MS) for the quantification of lanosterol and ergosterol**

UPLC-MS/MS was employed to quantify lanosterol and ergosterol in fungi cell samples. Samples were analyzed using freshly prepared calibration and quality control standards following the U.S. Food and Drug Administration bioanalytical method validation guidelines *(76)*. Samples were diluted by the addition of methanol, followed by sonication for 15 min, and gentle vortex mixing. Quantification was performed using freshly prepared analytical calibration and quality control standards, as matrix-matched calibration could not be employed due to the endogenous nature of both analytes. These samples (25 µL) were further processed with methanol (75 µL) containing internal standard (IS, 2 µg/mL) and formic acid (0.05 %, v/v). After vortexing the mixture, samples were filtered through a 0.45 µm 96-well Solvinert filter plate (Millipore, St. Louis, MO), centrifuged at 2000 rpm for 5 min at 4 °C, and injected into the Waters Acquity I coupled with Xevo® TQ-S mass spectrometer (UPLC-MS/MS). The chromatographic separation was achieved using a linear gradient elution program using a Waters Acquity UPLC BEH $C_{18}$ column (1.7 µm, 2.1 x 100 mm) with a VanGuard pre-column of a similar chemistry (Waters Co, Milford, MA), with flow rate of 0.4 mL/min. The aqueous phase comprised 0.1% formic acid in water (A) and the organic phase consisted of 0.1% formic acid in methanol (B). The gradient program was initiated at 20% mobile phase B from 0 to 0.5 min, increase to 98% B between 0.5 and 1.0 min. It was held at 98% B until 4.0 min, after which it was back to 20% B from 4.1 to 5.0 min to allow column equilibration. The column oven was set to 40 °C, while the autosampler was maintained at 10 °C. A strong needle wash solution composed of water, acetonitrile, methanol, and isopropyl alcohol (1:1:1:1, v/v) containing 0.1% formic acid, whereas the weak needle wash consisted of water, acetonitrile, and methanol (2:1:1, v/v) with 0.1% acetic acid. Atmospheric pressure chemical ionization (APCI) operated in positive multiple reaction monitoring (MRM) mode was used for the quantification of lanosterol and ergosterol. The APCI source temperature was maintained at 350 ºC, and the corona current was set to 3 µA. Data acquisition was accomplished using MassLynx software (version 4.2, Waters Corporation) and quantification was carried out using TargetLynx software. The optimized compound-specific parameters for UPLC–MS/MS analysis are as follows.

**Table S4. Compound parameters for quantification of analytes using UPLC-MS/MS**

| Analytes | Parent (*m/z*) | Daughter (*m/z*) | Cone (V) | Collision V) |
|---|---|---|---|---|
| Lanosterol | 409.3 | 109.1 | 24 | 32 |
| Ergosterol | 379.3 | 69.0 | 32 | 30 |

| | | | | |
|---|---|---|---|---|
| 6-methylprednisolone (Internal standard, IS) | 357.1 | 135.0 | 72 | 22 |

**Imaging mass spectrometry**

**Elemental Imaging**

Kidney sections were analyzed via laser ablation inductively coupled plasma of flight mass spectrometry (LA-ICP-TOF-MS) at the Biomedical National Elemental Imaging Resource (BNEIR) using an imageBIO266 laser ablation (LA) instrument (Element Scientific Lasers, Omaha, NE) interfaced to a Nu Instruments Vitesse (Wrexham, UK) ICP-TOF-MS via a Direct Concentric Injector. Laser ablation used a solid state 266 nm laser, which couples with biological materials but not glass slides. The laser was operated at a fluence of 3.5 J $cm^{-2}$ with a 250 Hz laser frequency. Images used a 2 μm square spot. The ICP-TOF-MS was tuned to operate with individual laser pulse widths of 5-6 milliseconds, using a nebulizer flow of 1150 ml $min^{-1}$ and helium flow through the chamber and imaging cup of 250 and 200-250 ml $min^{-1}$ respectively. The ICP-TOF-MS uses a segmented reaction cell with $H_2$ flow of 4 ml/min and He at a flow of 13 ml/min. The mass range monitored was from phosphorus at mass 31 to selenium at mass 80. Instrumentation was tuned daily to optimize response sensitivity and laser single pulse rate.

**Quantification**

LA and ICP-TOF-MS data were unified using Iolite software *(78),* which also performs background correction (using the signal from a gas-only blank between ablation for each line using a linear spline function) and converts raw counts into counts per second. Counts per second data were quantified into parts per million (ppm) using an in-house gel standard calibration series *(79)* consisting of six concentrations of elements of interest, ranging from 0 to 4000 femtograms total per spot. 3 g $L^{-1}$ gallic acid was added to increase the absorbance of gelatine at 266 nm to ensure similar ablation properties to the specimen.  Data reduction was performed in the Iolite 4 software package using the 3D Trace elements scheme. Gel standards were ablated before and after every specimen *(80)*. In addition, an in-house secondary--check standard consisting of 10 μm thick cryo-sectioned, homogenized bovine heart tissue was analyzed with every specimen. Conservative detection limits were established as three times the standard deviation of the mean of a user-defined sample-free region of the ablated material, and data shown are limit of detection (LOD) filtered, where values below the calculated LOD are removed from the image by assigning it missing value identifier.

### Post-processing analysis

Elemental images were processed using both *ScaleBarOn*, a custom Python-based tool developed for comparative scaling of multiple elemental image and automated composite elemental image generation and in *Muad'Data,* a custom Python-based tool developed for users of BNEIR to explore and save single-specimen data, either as individual elements or a multiple element overlays. LOD calculation and filtering as well as three element overlays were conducted in *Muad'Data.*
